# NeuralPlayground: A Standardised Environment for Evaluating Models of Hippocampus and Entorhinal Cortex

**DOI:** 10.1101/2024.03.06.583699

**Authors:** Clémentine C. J. Dominé, Rodrigo Carrasco-Davis, Luke Hollingsworth, Nikoloz Sirmpilatze, Adam L. Tyson, Devon Jarvis, Caswell Barry, Andrew M. Saxe

## Abstract

Neural processes in the hippocampus and entorhinal cortex are thought to be crucial for spatial cognition. A growing variety of theoretical models have been proposed to capture the rich neural and behavioral phenomena associated with these circuits. However, systematic comparison of these theories, both against each other and against empirical data, remains challenging. To address this gap, we present NeuralPlayground, an open-source standardised software framework for comparisons between theory and experiment in the domain of spatial cognition. This Python software package offers a reproducible way to compare models against a centralised library of published experimental results, including neural recordings and animal behavior. The framework implements three `Agents` embodying different computational models; three `Experiments` comprising publicly available neural and behavioral datasets; a customisable 2-dimensional `Arena` (continuous and discrete) able to generate common and novel spatial layouts; and a `Comparison` tool that facilitates systematic comparisons between models and data. Each module can also be used separately, allowing standardised and flexible access to influential models and data sets. We hope NeuralPlayground, available on GitHub^3^, provides a starting point for a shared, standardized, open, and reproducible computational understanding of the role of the hippocampus and entorhinal cortex in spatial cognition.

## 1. Introduction

Upon acquiring a novel experimental dataset, the route into data modeling, integrating the data with existing theories and performing comparative analyses is intricate. Beginning with data organization and visualization, the process progresses to creating a simulation environment for agent modeling and training. This might be followed by an in-depth hyperparameter tuning, then visualizing and quantifying results for robust comparisons. This iterative method is essential to compare new findings with the vast body of literature on the hippocampus and entorhinal cortex [1–3]. Indeed, an ideal approach might involve scrutinizing each experiment against every relevant theory. However, each new dataset or agent, with its unique format and challenges, adds to the workload significantly. The increasing volume of data and the breadth of phenomena under study further complicate this task [4–7].

To address the considerable challenges posed by this task, we introduce NeuralPlayground, an open-source, standardized framework designed to streamline the process of comparing entorhinal and hippocampal circuit (EHC) models and experimental data. The present version of the framework comprises three agents, including a successor representation model [8], an excitatory/inhibitory plasticity model [9], and the Tolman-Eichenbaum machine (TEM) [10]. It implements a customizable 2D arena (both continuous and discrete) that can replicate common experimental settings, such as T-maze, circular, and dynamic arenas. Finally, it facilitates simulations of the agents’ interactions in these environments, and contrasts these with experimental results. Presently, NeuralPlayground provides three such publicly available neural and behavioral datasets which have been cleaned and standardized to ensure accessibility [4–6]. Crucially, the framework provides a `Comparison` tool that seamlessly allows users to compare the results of artificial agents and experimental measurements from this array of `Agents`, `Arenas`, `Experiments`, and `Metrics`. For instance, upon introducing a new experiment, NeuralPlayground automates the comparison process with all implemented theories, expediting the critical task of theory-data integration (schematized in Fig. 1).

**Figure 1.**
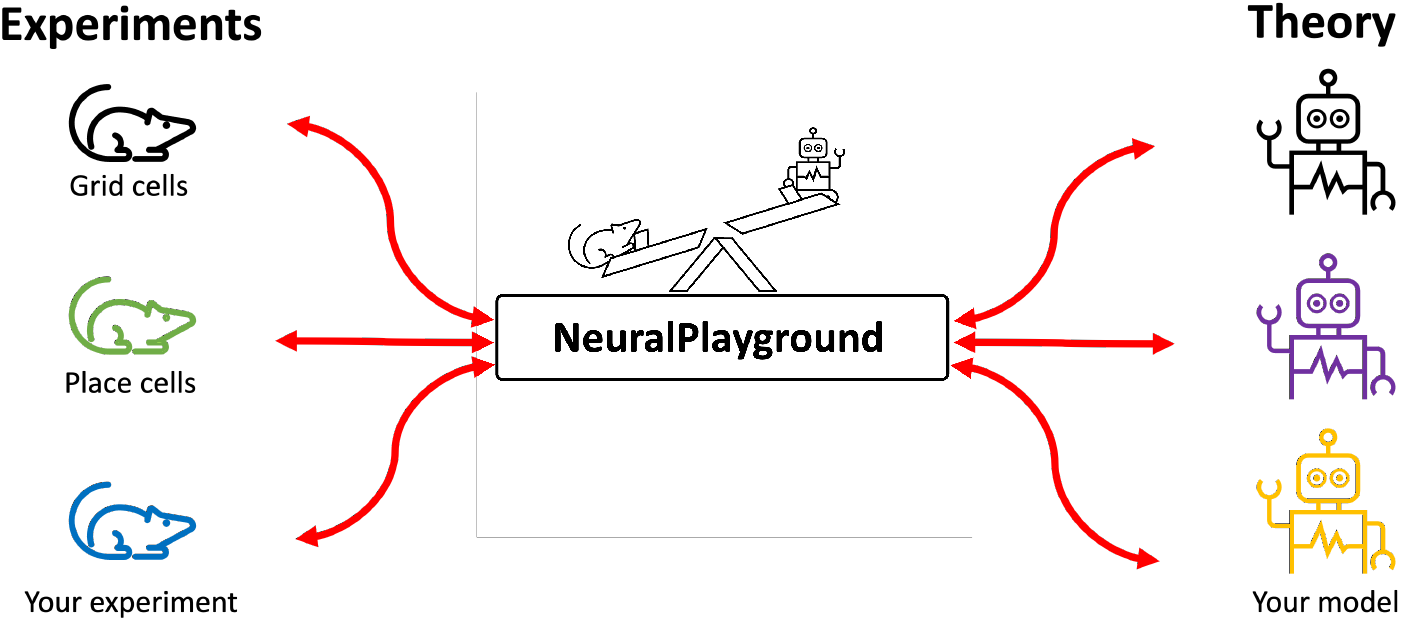
NeuralPlayground: An open-source standardized framework to compare hippocampus and entorhinal cortex models, aiding the loop between experiments and theoretical research in the field. Upon incorporating a new experiment, NeuralPlayground automates the comparison process with all implemented theories (rightward red arrow). Conversely, upon implementing a new theory embodied as a software agent, NeuralPlayground automates the comparison to all experiments (leftward red arrow) and places these comparisons alongside other theories.

## 2. Motivation

### 2.1. The Hippocampus and Entorhinal Cortex

NeuralPlayground focuses on the role of hippocampus and entorhinal cortex in spatial cognition. The selection of these particular brain structures and function reflects the substantial amount of empirical and theoretical research associated with them [1]. Moreover, these circuits contain diverse functional cell types characterised by firing properties, including place cells [11], grid cells [12], and head direction cells [13]. Consequently, these regions provide a clear link between neural phenomena and cognitive or behavioural outcomes which can be modelled theoretically. Specifically, at the population or circuit level, the hippocampus and entorhinal cortex has been proposed to encode a “cognitive map” [11, 14, 15]. This enables mammals to efficiently and flexibly navigate the environment. The representations of space encoded by this cognitive map have been extensively studied, both experimentally [8–10, 12, 13, 16–20] and theoretically [8–10, 21–23]; see [1–3] for a review. A variety of models drawing on different principles such as attractor networks [14, 23, 24], spiking neuron models [25], and reinforcement learning [8, 26] have similarly been discovered. This affords extensive comparison with experiments and has sparked new research questions and a better understanding of navigation and learning mechanisms. Thus, in sum, hippocampus and entorhinal cortex when employed in spatial cognition have distinct measurable neural mechanisms that are observable in behavior (both organic and artificial). The resulting abundance of experimental and theoretical data covers various task settings. However, this abundance necessitates software for data management and comparison, to fully harness the potential insights. NeuralPlayground is designed to meet this need, offering a solution for navigating and maximizing the value of this extensive data.

### 2.2. Standardised Framework

While the importance of interaction between experiments and theory is clear, the available tools facilitating comparison are limited. The first challenge for the field in building a reliable comparison comes from **1) the availability and accessibility of the data in a standard, labeled format**. Even though the field is pushing forward in the direction of more open source collaboration, it remains difficult to parse many task details and access behavioral and neural data. Furthermore, users are confronted by a variety of data formats and pre-processing steps performed by each study, with disparate documentation of these methods. As a result, new modelling work typically only compares to a fraction of the available evidence and paradigms. A second challenge comes from the fact that **2) models and experiments are not described with the same focus, detail or levels of abstraction**. For example, not all models are able to generate neural data/behavior or interact with a task. Identifying which regimes each model is valid for, and which phenomena are relevant to compare against is not always straightforward. On the experimental end, the lack of descriptors of experimental environments that can be used in models could prevent accurate computational task modeling. Another prevalent issue is **3) the lack of standard or easy ways for models to interact with the task**. To test models, researchers often generate bespoke simulations of animal behavior and interactions with an environment, hence duplicating work, introducing variability, and increasing chances of errors in re-implementations. Finally, **4) there is no consensus on standard ways to compare model predictions with empirical data**. Altogether, this ultimately leads to a lack of standardisation and makes the aim of general accessibility challenging.

A better understanding of the ways models behave in different situations, beyond those reported in their respective publications, will lead to novel experimental research and illuminate the pathologies and shortcomings of each model. We aim to facilitate this by providing a tool for a simple inclusion of new models, experiments, and comparison methods. In many other areas, achieving scale has been critical to advances in the field. We aim to facilitate the scaling of model comparison which we hope will further aid cooperation and collaboration in the field.

## 3. Previous Work

Within hippocampus and entorhinal cortex, previous work such as RatInABox [27] and Neuro-Nav [28] have successfully paved the way in building reproducible frameworks for neuroscience. Neuro-Nav and RatInABox share a similar overarching goal but emphasize distinct aspects of this endeavor. RatInABox complements our approach by allowing a wider variety of realistic animal trajectories and associated neural data, for cells that are spatially and/or velocity selective in complex, continuous surroundings. Neuro-Nav is an open-source platform for reinforcement learning (RL) that is based on the principles of neuroscience. It provides a collection of standard environments and a large library of RL algorithms inspired by classical studies in rodents and humans, and focuses on behaviour. In contrast to both, NeuralPlayground implements a broad class of models which span RL, cognitive connectionist/deep learning models, and systems neuroscience circuit models. Furthermore, we allow for both continuous and discrete action policies and environments with a particular focus on comparisons between experimental and model neural responses in these environments.

The idea of a benchmark has been successfully implemented for models of the visual system [29] and computational cognitive neuroscience models [30]. These benchmarks are now widely used by the community as a baseline for testing models and have prompted novel research. Our benchmark methodology differs from the standard approach because of the wide variation in environmental conditions reported in the EHC literature – across both experiments and models – makes defining a single conclusive performance metric challenging. Instead, we have adopted an automated approach that generates a range of metrics and visual representations. This strategy enables a comprehensive comparative analysis, both quantitative and qualitative, of outcomes from model simulations and experimental results.

## 4. NeuralPlayground library

### 4.1. NeuralPlayground Modules

The NeuralPlayground software package, built in Python, is composed of four main components organised in four modules. Each module is built in a similar fashion: A core class defines a shared set of methods. Each instantiation of a new agent, environment, experimental dataset or metric can be easily implemented as a child class, inheriting these methods. We now describe each of these modules in turn.

#### Agent

The `Agent` class includes a set of functions that control the way intelligent systems interact with their environment. An `Agent` receives observations from the environment (reward, visual cues, etc.) and uses these to select an action, which in turn will update both its state and the state of the environment, generating new observations. More generally, the `Agent` can be thought of as an animal performing the task in the simulated `Experiment`. We have de novo implementations of three influential models of the hippocampus and entorhinal cortex, namely the successor representation approach of Stachenfeld et al., (2017) [8], the excitatory/inhibitory plasticity model of Weber & Sprekeler, (2018) [9], and the Tolman-Eichenbaum Machine (TEM) of Whittington et al., (2020) [10]. These models are summarized in Fig. 2 and span a broad class of models including RL, systems neuroscience circuit models, and cognitive connectionist/deep learning models.

**Figure 2.**
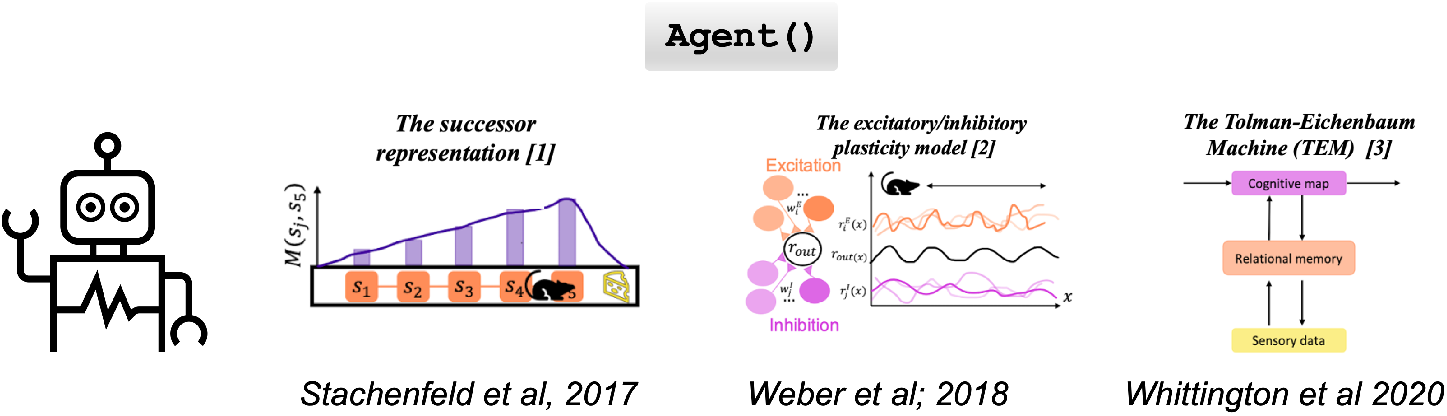
Simplified schematics of the `Agents` implemented. (Left) The successor representation (SR) model which represents the likelihood of moving into a state *s*_*j*_ from a previous state *si* in the SR matrix *M* (*s*_*i*_, *s*_*j*_) [8]. This model has been used to show that place fields skew backward toward previous states when the value function is flexibly learned independently from the SR matrix *M* (*s*_*i*_, *s*_*j*_). (Middle) The Oscillatory-Interference model [9] on a linear track. A threshold-linear model (*r*_*out*_) builds a grid-like spatial receptive field from inputs received from spatially tuned excitatory (orange) and inhibitory (pink) neurons (*r*_*i*_ and *r*_*j*_ respectively). (Right) Depiction of the Tolman-Eichenbaum Machine (TEM) at one time point: The model takes inspiration from Tolman’s theory of an internal cognitive map (pink) [14], and combines it with sensory data (yellow) via the relational memory (orange) of Eichenbaum [10, 31]

**Figure 3.**
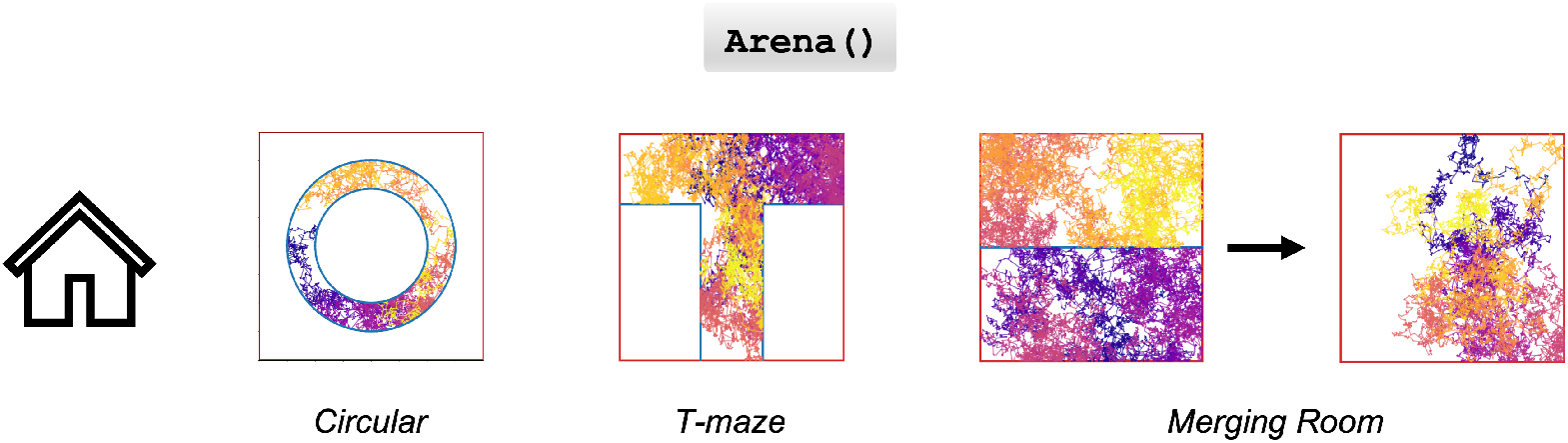
Illustration of various `Arena` layouts commonly used in hippocampus and entorhinal cortex experiments, showcasing an `Agent` employing a random walk strategy in distinct environments: Circular, T-Maze, and Merging Room. In the Merging Room scenario, the arena initially features a dividing wall which is subsequently removed, demonstrating the dynamic adaptability of the environment.

#### Arena

The `Arena` module creates a space within which an `Agent` can navigate, learn, and interact, mirroring the physical configurations of experimental setups used in behavioral and neural data collection. It enables the construction of any two-dimensional layout, both discrete and continuous, using walls as building elements. This flexibility allows for the creation of intricate experimental designs, such as interconnected rooms, T-mazes, or circular tracks, as depicted in Fig.3. Additionally, dynamic `Arenas` that change over time, like the merging room experiment from Wernle et al. (2018), can be accommodated (Fig.3), offering a versatile platform for a wide range of experimental replications and explorations.

#### Experiments

The `Experiments` module facilitates access to a curated collection of open-source experimental data, including neural recordings and behavioral observations, and a suite of plotting functions and visualization tools (Fig. 4). The primary objective of this module within our package is to establish a centralized and standardized portal for accessing pertinent experimental data in the field. This data is pre-processed and annotated for ease of use, streamlining the exploration and analysis process. Each data set is organised into recording sessions with an attributed recording number (`rec index`), given as a list at the initialisation of the class. Our package allows for versatile visualizations and data access for experimental results. One can plot the activity of a specific tetrode using `.plot_recording_tetr(index)` (see red-blue heatmaps in Fig. 4), visualize the movement trajectory within the `Arena` with `.plot_trajectory(index)` (shown in Fig. 4 with the pink-yellow line plots), and retrieve detailed experimental information using `.show_keys()`. We have implemented three data sets: (1) Sargolini et al. (2006) [6], in which tetrode recordings were made as rats freely explored a flat square environment and found that, while Layer II of MEC was predominantly composed of grid cells, deeper layers of MEC have both grid, head direction and conjunctive grid *×* head direction cells. (2) Hafting et al. (2008) [5], which examines a linear environment and found that the phase precession of firing patterns (cells fire out of phase with the ambient theta wave pattern in the surrounding areas) primarily in Layer II of MEC drives phase precession of place cells in hippocampus. (3) Wernle et al. (2018) [4], which considers a dynamic environment of two partitioned rooms which have individual grid-cell representations. Partway through a session the environment changes and the partition is remove - merging the rooms, resulting in the rapid reorganisation of the grid cells such that the periodicity of previously connected spaces remains but now with new consistent periodicity also established where the partition used to be.

**Figure 4.**
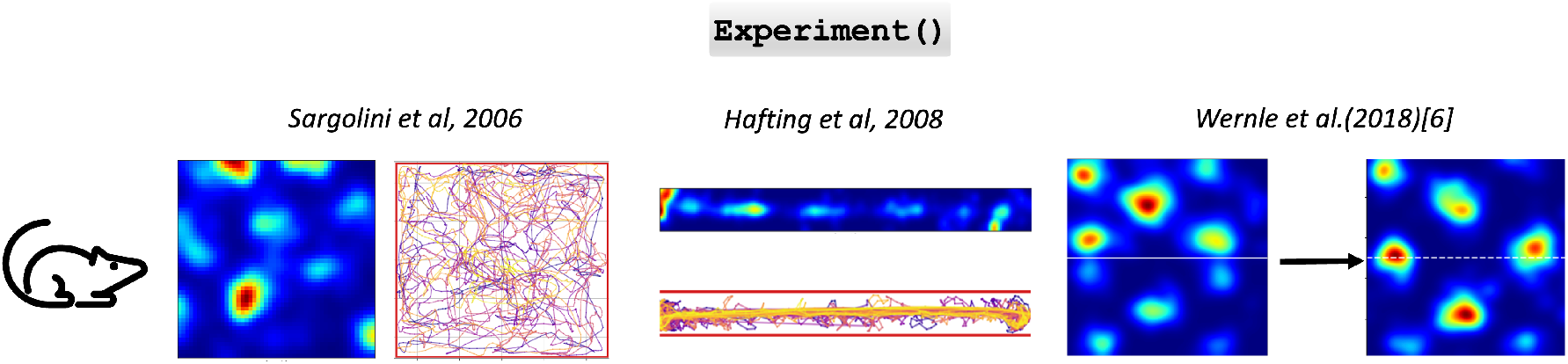
Sample data from the `Experiment` modules. Grid like rate map and position of the rat in the environment from the Sargolini et al. [6], Hafting et al. [5] and Wernle et al. [4] experimental results respectively.

#### Comparison

Finally, the `Comparison` tool is a versatile feature designed to enhance user experience and research efficiency. It grants users the flexibility to select from a range of `Agents`, `Arenas`, and `Experiments`, and to determine specific plots they wish to generate. This functionality not only simplifies the process of creating visual representations but also enables a comprehensive and systematic comparison across all integrated `Agents` and `Experiments`. By offering customizable comparison parameters, this tool facilitates a deeper analysis and understanding of the interactions and outcomes between various agents and experimental settings.

### 4.2. Module Use

The framework allows for flexible access to theoretical and experimental paradigms. Each module can be used separately, to easily explore and analyze experimental data and better understand, test or modify any of the implemented models. Additionally, different `Arenas` can be initialised with custom wall structures, or following the spatial design of real-life experiments. We provide examples of module instantiation in detailed jupyter notebook *examples*.^4^

The `Agents` and `Arenas` within the framework interact similarly to those found in the OpenAI Gym environment [32]. Each `Agent` within this framework receives a stream of observations from the surrounding environment, including rewards, visual cues, and other relevant information. These observations inform the `Agent`’s decision-making process, culminating in the execution of an action. The execution of actions, in turn, leads to updates in the `Agent`’s internal state and simultaneously updates the state of the environment. Consequently, this dynamic interaction generates new observations (Fig. 5).

**Figure 5.**
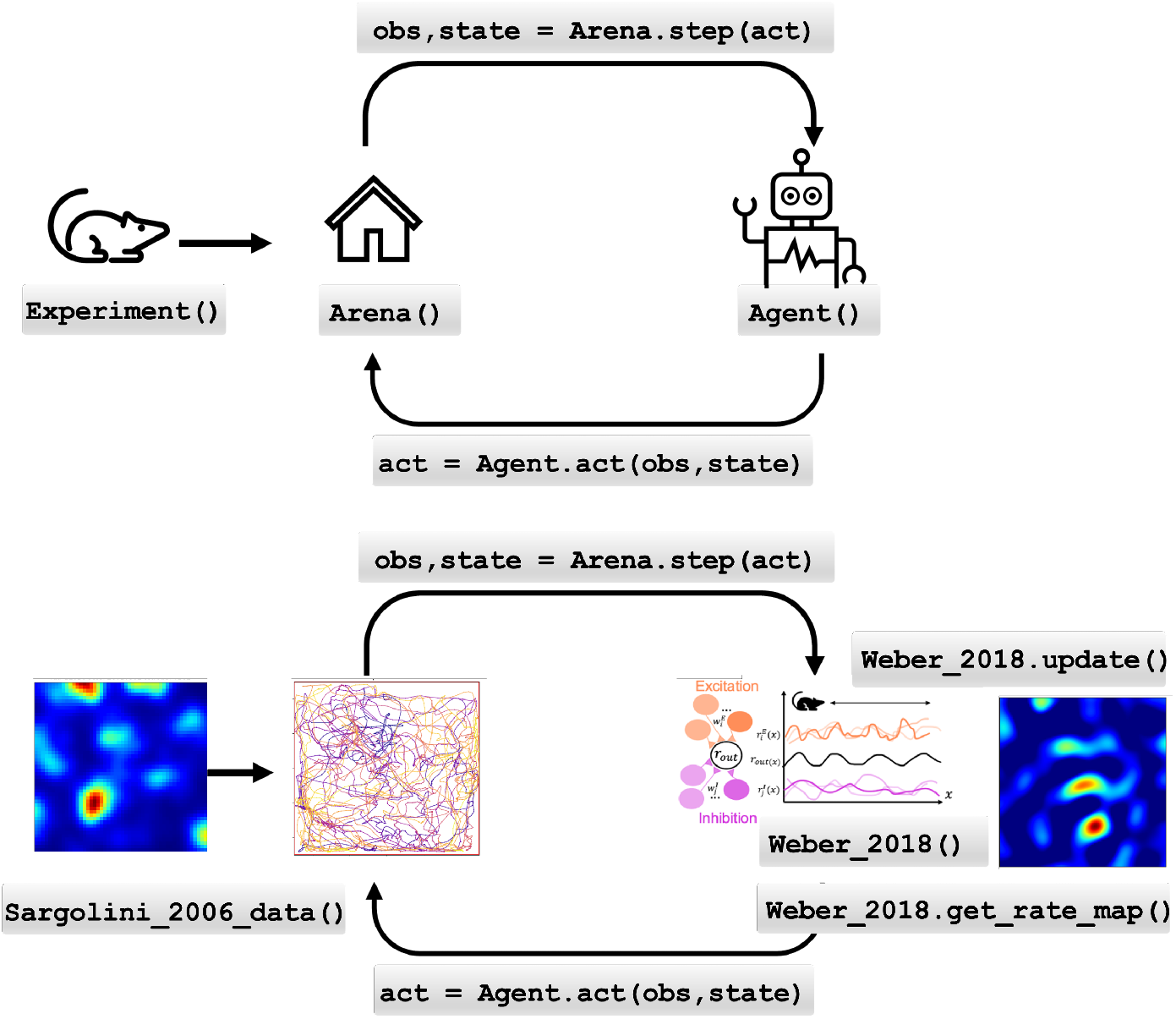
Illustration depicting a practical application of the NeuralPlayground framework. Top: Schematic of the modules’ interactions: The greyed boxes specify the lines of code performing the interaction represented by black arrows. The `Agent` obtains feedback from the environment, such as rewards and visual information, and utilizes this information to make decisions that result in changes to its own state and the state of the environment, leading to the generation of fresh observations. Moreover, the data extracted from the `Experiment` class can be employed to construct `Arenas` that replicate elements of the experimental setup.. Bottom: For a concrete setting, we select the `Weber_2018 Agent`, the `Sargolini_2006_data Experiment`, and the `Weber_2018 Arena` class generated by the `Sargolini_2006_data Experiment` class. As the model walks through the experimental environment following the animal’s path, continually updating its neural representation, it ultimately generates grid-like representations, as visualised with the function `.get_rate_map()`.

Furthermore, the data extracted from the `Experiment` class can be effectively leveraged to construct `Arena` configurations that resemble key elements of the experimental setups. For example, the user can set the `Agent` to follow the behavioral trajectory of the animal recorded during the experiment, or use a built in policy to move around if available.

The `Agent` example jupyter notebooks^5^ provide examples where one can create this interaction with minimal code. These begin by initializing an `Agent` and `Arena` of choice, which are built with additional specific methods, such as `.get_neural_response()` and `.get_rate_map()` for `Agents`, and `.plot_experimental_results()` for `Arenas` (Fig. 5). The interaction between agent and environment can then be instantiated as a `for` loop, as shown in Code List. 1 which is used with the `agent.get_rate_map()` function:

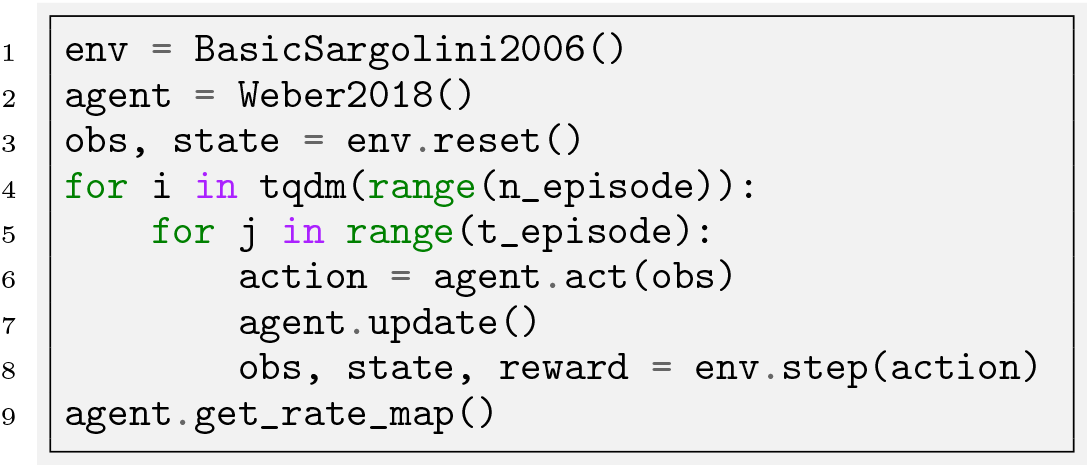

Code Listing 1: Example interaction between an agent, namely the excitatory/inhibitory plasticity model of Weber & Sprekeler (2018) [9], and an environment, the standard square room from Sargolini et al. (2006) [6].

The output of this `Agent` and `Arena` interaction is summarised in Fig. 5. The initialisation of the `Arenas` and `Agents` are automated in the background and the results are displayed for qualitative comparison (Fig. 6). Because of the range of phenomena and levels of detail that have been addressed by prior experiments and models, NeuralPlayground does not implement one global leaderboard metric. Instead, we provide a tool for systematic comparison to the evidence in the field using the metrics of interest to the user. Thus, NeuralPlayground will support diverse qualitative and quantitative comparisons as the field progresses. We show an example use case in the *comparison_examples*^6^.

**Figure 6.**
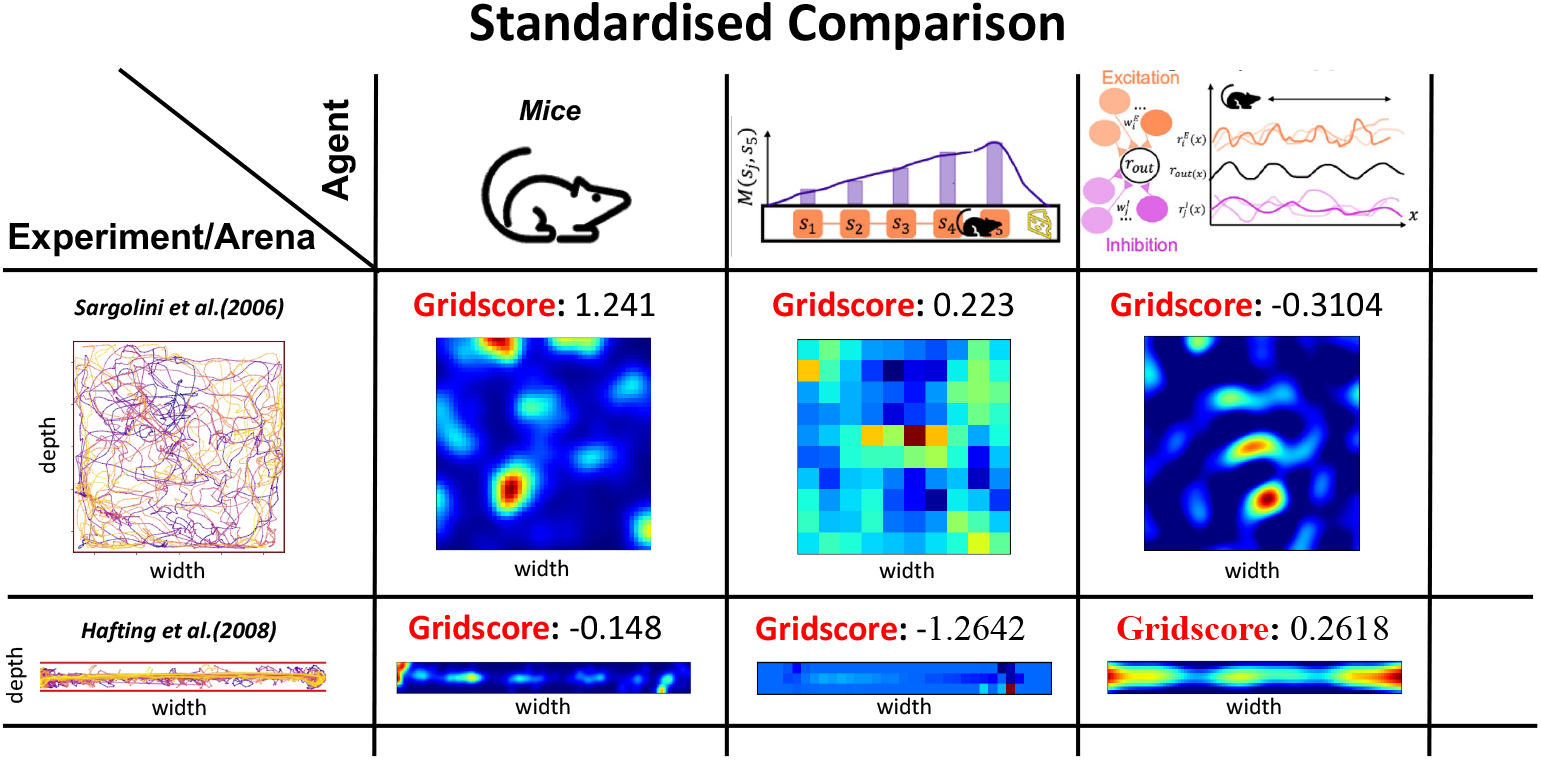
The `Comparison Board` automatically generates and displays results from various experimental runs. In this organizational layout, each row corresponds to a distinct experimental `Arena`, while each column represents a specific `Agent`. This example displays the output from using two `Agents`, namely the SR model developed by Stachenfeld and the Weber model, and two `Environments`, namely the Sargolini and Hafting experiments. The comparison board deploys each agent in both the Sargolini and Hafting environments, enabling a comparative analysis of their performance in relation to the neural representations observed in the experimental data from mice. As a qualitative comparison, here the rate map of one neuron is visualised. As a quantitative comparison, we calculate the gridness score, as defined by Sargolini et al. in 2006.

To demonstrate the available analyses from this tool we provide example outputs in Figs. 6 and 7. We assess the performance of the `Agent` against a set of selected experimental observations, roughly categorized as qualitative and quantitative evidence available within the `Comparison` module.

**Figure 7.**
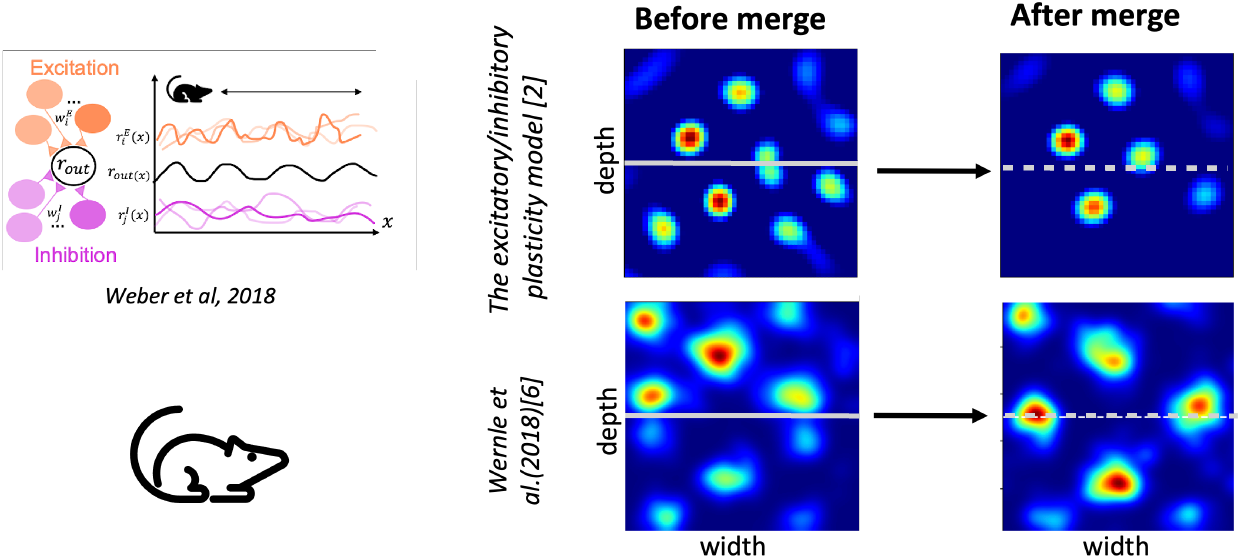
Example qualitative comparison in the merging rooms experiment. The merging room experiment involves allowing the agent to navigate an initially partitioned environment, with subsequent removal of the barrier. As documented experimentally in [4], neural responses merge at the boundary. This qualitative behaviour is successfully reproduced by the Weber model [9]

#### Qualitative analysis

NeuralPlayground can generate response rate maps for model neurons, averaged over periods of interaction with the environment (Fig. 6. These can be used to identify the presence of different types of cells including Place Cells [11], Grid Cells [12], Boundary Vector Cells [33], Border Cells [34], and Object Vector Cells [35]. Another example of qualitative evidence comes from the merging rooms experiment [4, 9], which examines how representations change when a wall is removed from an environment, as exemplified in Fig. 7.

#### Quantitative analysis

In accordance with the methodology established by Sargolini et al. [6], NeuralPlayground can calculate a quantitative metric known as the “gridness score”, ranging from -2 to 2, with large magnitude scores indicating grid-like responses (Fig. 6). Additional scoring metrics can be easily and systematically generated by writing a new metric class that may be relevant to the research context.

## 5 Open-source collaborative framework

The NeuralPlayground open-source software was built to be a collaborative and lasting project. The code is freely available online under an *MIT license*^7^. All contributions to the repository are acknowledged through the all-contributors bot^8^. To contribute, refer to the “Documents” section in the repository. Contributions to a module can be made in two main ways: by creating a new class or by adding methods to an existing one.

Long term software support and maintenance is provided by the Gatsby/SWC neuroinformatics team. This includes library maintenance, test implementation, and the introduction of bots to streamline workflows, among other responsibilities. Additionally, we adhere to reproducible, inclusive, and collaborative project design guidelines, as outlined in [36]. We also encourage users to follow our community’s recommended guidelines and code of conduct^9^.

Collectively, we hope this framework will aid and build the community of researchers who seek to advance our understanding of computational mechanisms within the hippocampus, entorhinal cortex, and other brain regions. We suggest several ways to begin building upon the NeuralPlayground Framework:

- `Arena`: Primarily, the expansion of the `Arena` module will include a new sub-class for 3D `Arenas` and sensory stimuli inspired by OpenAIGym [32] or Deepmind lab [37].
- `Experiments`: Broaden the repository of implemented experimental datasets pertinent to the hippocampus and entorhinal cortex research, focusing on publicly available sources ([38–40]). The true potential of the software will be realized as it incorporates an increasingly diverse array of experimental datasets and comparison metrics. This expansion will not only enrich the software’s capabilities but also foster a more comprehensive and nuanced understanding within the field.
- `Agent`: One can also contribute by adding a new `Agent` [41, 42] or extending one that is already implemented. The package was designed to allow versatile classes of `Agents`, and the addition of behavior and replay models [43].
- `Comparison`: One way to contribute to the `Comparison` is by adding new metrics of comparison such as grid score.

## 6. Discussion

In the era of large datasets and increasingly capable computational models, it is critical to facilitate diverse comparisons between theory and experiment. In the hippocampus and entorhinal cortex, prior research, exemplified by RatInABox [27] and Neuro-Nav [28], has effectively laid the groundwork for constructing replication-centric frameworks in neuroscience. Another key ingredient in managing the thorough comparison of datasets and models has proven to be open collaborative community efforts such as BrainScore [44] and the CCNlab benchmark [30] which have fostered rapid progress in other areas of neuroscience such as visual perception.

Here we release a beta version of the NeuralPlayground framework, with three `Agents` implemented from established computational models; three data sets from previous `Experiments`; and with a customizable 2-Dimensional `Arena`. The software allows for a diversity of use cases, including aiding data analyses, and allowing evaluation of models beyond their initial scope in environment, metric or comparison to relevant experimental observations. Because of the range of computations that the hippocampus and entorhinal cortex have been proposed to support, and the correspondingly diverse set of models used to account for aspects of this data, NeuralPlayground does not implement one summary benchmark metric but provides an expandable suit of benchmarks through the `Comparison` module. Primarily, the NeuralPlayground open-source software package centralizes and facilitates access and testing of models against experimental data. The outcomes we hope will arise from this effort are threefold. Firstly, NeuralPlayground will lead to a standardization of models and experimental methods for charaterizing neural responses in the EHC. Secondly, it will facilitate the creation of many quantitative and qualitative comparisons of value to the community. And lastly, a long-lasting community that guides, contributes to and nurtures this software will grow and help build a virtuous loop between experimentalists and theorists.

## 7. Acknowledgements

This research was funded in whole, or in part, by the Wellcome Trust [216386/Z/19/Z]. C.D., R.C-D, L.H, was supported by the Gatsby Charitable Foundation (GAT3755). D.J. is a Google PhD Fellow and Commonwealth Scholar. Further, A.S. was supported by the Sainsbury Wellcome Centre Core Grant (219627/Z/19/Z) and A.S. is a CIFAR Azrieli Global Scholar in the Learning in Machines & Brains program. C.B. was funded by the Wellcome Trust 212281/Z/18/Z.

## Footnotes

3 github.com/SainsburyWellcomeCentre/NeuralPlayground

4 github.com/NeuralPlayground/Jupyter_notebooks

5 github.com/NeuralPlayground/Agent_jupyter_notebooks

6 github.com/NeuralPlayground/comparisons_examples/

7 github.com/NeuralPlayground/documents

8 github.com/NeuralPlayground/all-contributors

9 github.com/NeuralPlayground/documents

## References

[1] Edvard I Moser, May-Britt Moser, and Bruce L McNaughton. Spatial representation in the hippocampal formation: a history. Nature neuroscience, 20(11):1448–1464, 2017.

[2] James CR Whittington, David McCaffary, Jacob JW Bakermans, and Timothy EJ Behrens. How to build a cognitive map: insights from models of the hippocampal formation. arXiv preprint arXiv:2202.01682, 2022.

[3] György Buzsáki and David Tingley. Space and time: the hippocampus as a sequence generator. Trends in cognitive sciences, 22(10):853–869, 2018.

[4] Tanja Wernle, Torgeir Waaga, Maria Mørreaunet, Alessandro Treves, May-Britt Moser, and Edvard I Moser. Integration of grid maps in merged environments. Nature neuroscience, 21(1):92–101, 2018.

[5] Torkel Hafting, Marianne Fyhn, Tora Bonnevie, May-Britt Moser, and Edvard I Moser. Hippocampus-independent phase precession in entorhinal grid cells. Nature, 453(7199):1248–1252, 2008.

[6] Francesca Sargolini, Marianne Fyhn, Torkel Hafting, Bruce L McNaughton, Menno P Witter, May-Britt Moser, and Edvard I Moser. Conjunctive representation of position, direction, and velocity in entorhinal cortex. Science, 312(5774):758–762, 2006.

[7] Guifen Chen, John Andrew King, Yi Lu, Francesca Cacucci, and Neil Burgess. Spatial cell firing during virtual navigation of open arenas by head-restrained mice. eLife, 7:e34789, jun 2018. ISSN 2050-084X. doi: 10.7554/eLife.34789. URL 10.7554/eLife.34789.

[8] Kimberly L Stachenfeld, Matthew M Botvinick, and Samuel J Gershman. The hippocampus as a predictive map. Nature neuroscience, 20(11):1643–1653, 2017.

[9] Simon Nikolaus Weber and Henning Sprekeler. Learning place cells, grid cells and invariances with excitatory and inhibitory plasticity. Elife, 7:e34560, 2018.

[10] James CR Whittington, Timothy H Muller, Shirley Mark, Guifen Chen, Caswell Barry, Neil Burgess, and Timothy EJ Behrens. The tolman-eichenbaum machine: Unifying space and relational memory through generalization in the hippocampal formation. Cell, 183(5):1249–1263, 2020.

[11] John O Keefe and Lynn Nadel. The hippocampus as a cognitive map. Clarendon Press, 1978.

[12] Marianne Fyhn, Torkel Hafting, Menno P Witter, Edvard I Moser, and May-Britt Moser. Grid cells in mice. Hippocampus, 18(12):1230–1238, 2008.

[13] Neil Burgess, Francesca Cacucci, Colin Lever, and John O’keefe. Characterizing multiple independent behavioral correlates of cell firing in freely moving animals. Hippocampus, 15(2):149–153, 2005.

[14] Edward C Tolman. Cognitive maps in rats and men. Psychological review, 55(4):189, 1948.

[15] RU Muller, JL Kubie, EM Bostock, JS Taube, and GJ Quirk. Brain and space, 1991.

[16] Elizabeth Bostock, Robert U Muller, and John L Kubie. Experience-dependent modifications of hippocampal place cell firing. Hippocampus, 1(2):193–205, 1991.

[17] Robert U Muller and John L Kubie. The effects of changes in the environment on the spatial firing of hippocampal complex-spike cells. Journal of Neuroscience, 7(7):1951–1968, 1987.

[18] John L Kubie and Robert U Muller. Multiple representations in the hippocampus. Hippocampus, 1(3):240–242, 1991.

[19] Hanne Stensola, Tor Stensola, Trygve Solstad, Kristian Frøland, May-Britt Moser, and Edvard I Moser. The entorhinal grid map is discretized. Nature, 492(7427):72–78, 2012.

[20] Emilio Kropff, James E Carmichael, May-Britt Moser, and Edvard I Moser. Speed cells in the medial entorhinal cortex. Nature, 523(7561):419–424, 2015.

[21] John O’keefe and Neil Burgess. Dual phase and rate coding in hippocampal place cells: theoretical significance and relationship to entorhinal grid cells. Hippocampus, 15(7):853–866, 2005.

[22] Lisa M Giocomo, May-Britt Moser, and Edvard I Moser. Computational models of grid cells. Neuron, 71(4): 589–603, 2011.

[23] Christopher J Cueva and Xue-Xin Wei. Emergence of grid-like representations by training recurrent neural networks to perform spatial localization. arXiv preprint arXiv:1803.07770, 2018.

[24] Edmund T Rolls. An attractor network in the hippocampus: theory and neurophysiology. Learning & memory, 14(11):714–731, 2007.

[25] Misha V Tsodyks, William E Skaggs, Terrence J Sejnowski, and Bruce L McNaughton. Population dynamics and theta rhythm phase precession of hippocampal place cell firing: a spiking neuron model. Hippocampus, 6 (3):271–280, 1996.

[26] Juliet Y Davidow, Karin Foerde, Adriana Galván, and Daphna Shohamy. An upside to reward sensitivity: the hippocampus supports enhanced reinforcement learning in adolescence. Neuron, 92(1):93–99, 2016.

[27] Tom M George, William de Cothi, Claudia Clopath, Kimberly Stachenfeld, and Caswell Barry. Ratinabox: A toolkit for modelling locomotion and neuronal activity in continuous environments. bioRxiv, pages 2022–08, 2022.

[28] Arthur Juliani, Samuel Barnett, Brandon Davis, Margaret Sereno, and Ida Momennejad. Neuro-nav: A library for neurally-plausible reinforcement learning. In The 5th Multidisciplinary Conference on Reinforcement Learning and Decision Making, 2022.

[29] Martin Schrimpf, Jonas Kubilius, Ha Hong, Najib J Majaj, Rishi Rajalingham, Elias B Issa, Kohitij Kar, Pouya Bashivan, Jonathan Prescott-Roy, Franziska Geiger, et al. Brain-score: Which artificial neural network for object recognition is most brain-like? BioRxiv, page 407007, 2020.

[30] Nikhil Xie Bhattasali, Momchil Tomov, and Samuel Gershman. Ccnlab: A benchmarking framework for computational cognitive neuroscience. In Thirty-fifth Conference on Neural Information Processing Systems Datasets and Benchmarks Track (Round 1), 2021.

[31] Howard Eichenbaum and Neal J Cohen. Can we reconcile the declarative memory and spatial navigation views on hippocampal function? Neuron, 83(4):764–770, 2014.

[32] Greg Brockman, Vicki Cheung, Ludwig Pettersson, Jonas Schneider, John Schulman, Jie Tang, and Wojciech Zaremba. Openai gym. arXiv preprint arXiv:1606.01540, 2016.

[33] Caswell Barry, Colin Lever, Robin Hayman, Tom Hartley, Stephen Burton, John O’Keefe, Kate Jeffery, and N Burgess. The boundary vector cell model of place cell firing and spatial memory. Reviews in the Neurosciences, 17(1-2):71–98, 2006.

[34] Trygve Solstad, Charlotte N Boccara, Emilio Kropff, May-Britt Moser, and Edvard I Moser. Representation of geometric borders in the entorhinal cortex. Science, 322(5909):1865–1868, 2008.

[35] Øyvind Arne Høydal, Emilie Ranheim Skytøen, Sebastian Ola Andersson, May-Britt Moser, and Edvard I Moser. Object-vector coding in the medial entorhinal cortex. Nature, 568(7752):400–404, 2019.

[36] The Turing Way Community. The Turing Way: A handbook for reproducible, ethical and collaborative research, July 2022. URL 10.5281/zenodo.6909298.

[37] Charles Beattie, Joel Z Leibo, Denis Teplyashin, Tom Ward, Marcus Wainwright, Heinrich Küttler, Andrew Lefrancq, Simon Green, Víctor Valdés, Amir Sadik, et al. Deepmind lab. arXiv preprint arXiv:1612.03801, 2016.

[38] Caswell Barry, Lin Lin Ginzberg, John O’Keefe, and Neil Burgess. Grid cell firing patterns signal environmental novelty by expansion. Proceedings of the National Academy of Sciences, 109(43):17687–17692, 2012.

[39] Chen Sun, Wannan Yang, Jared Martin, and Susumu Tonegawa. Hippocampal neurons represent events as transferable units of experience. Nature neuroscience, 23(5):651–663, 2020.

[40] Guifen Chen, John Andrew King, Yi Lu, Francesca Cacucci, and Neil Burgess. Spatial cell firing during virtual navigation of open arenas by head-restrained mice. Elife, 7, 2018.

[41] Christopher J. Cueva and Xue-Xin Wei. Emergence of grid-like representations by training recurrent neural networks to perform spatial localization. 2018. doi: 10.48550/ARXIV.1803.07770. URL https://arxiv.org/abs/1803.07770.

[42] Neil Burgess, Caswell Barry, and John O’keefe. An oscillatory interference model of grid cell firing. Hippocampus, 17(9):801–812, 2007.

[43] Jesse P Geerts, Fabian Chersi, Kimberly L Stachenfeld, and Neil Burgess. A general model of hippocampal and dorsal striatal learning and decision making. Proceedings of the National Academy of Sciences, 117(49): 31427–31437, 2020.

[44] Martin Schrimpf, Jonas Kubilius, Ha Hong, Najib J. Majaj, Rishi Rajalingham, Elias B. Issa, Kohitij Kar, Pouya Bashivan, Jonathan Prescott-Roy, Franziska Geiger, Kailyn Schmidt, Daniel L. K. Yamins, and James J. DiCarlo. Brain-score: Which artificial neural network for object recognition is most brain-like? bioRxiv preprint, 2018. URL https://www.biorxiv.org/content/10.1101/407007v2.

